# *Dmon1* and *rab7* interact to regulate glutamate receptor GluRIIA levels at the larval *Drosophila* neuromuscular junction

**DOI:** 10.1101/2020.03.21.999524

**Authors:** Anagha Basargekar, Shweta Yogi, Zeeshan Mushtaq, Senthil Deivasigamani, Vimlesh Kumar, Girish Ratnaparkhi, Anuradha Ratnaparkhi

## Abstract

Regulation of post-synaptic receptors plays an important role in determining synaptic strength and plasticity. The *Drosophila* larval neuromuscular junction (nmj) has been used extensively as a model to understand some of these processes. In this context, we are interested in the role of *Drosophila* Monensin sensitive protein 1 (DMon1) in regulating glutamate receptor (GluRIIA) levels at the nmj. *Dmon1* is an evolutionarily conserved protein which, in complex with CCZ1, regulates the conversion of early endosomes to late endosomes through recruitment of Rab7. C-terminal deletion mutants of *Dmon1* (*Dmon1*^*Δ181*^) exhibit lethality. The escapers have a short life span and exhibit severe motor defects. At the nmj, these mutants show a defects in synaptic morphology and a strong increase in glutamate receptor GluRIIA levels. The mechanism by which *Dmon1* regulates GluRIIA is unclear.

In this study, we have described the characterization the mutation in an EMS mutant referred to as *pog*^*1*^ and demonstrate this mutant to be an allele of *Dmon1*. Further, we have examined the role of *rab7* in regulation the of GluRIIA. We show that similar to *Dmon1*, knock-down of *rab7* using RNAi in neurons, and not muscles, leads to an increase in GluRIIA. Loss of one copy each of *Dmon1* with *rab7* leads to a synergistic increase in receptor expression. Further, overexpression of an activated Rab7 can rescue the GluRIIA phenotype observed in *Dmon1*^*Δ181*^ mutants. Together, these results highlight a neuronal role for Rab7 in GluRIIA regulation and underscores the important of the endo-lysosomal pathway in this process.

## Introduction

Post-synaptic neurotransmitter receptors play an important role in determining synaptic strength and plasticity. Receptor levels are altered in response to changes in neurotransmitter release to maintain homeostasis. The *Drosophila* larval neuromuscular junction (nmj) has been used for a long time as a model to study synaptic development and homeostasis. These synapses are glutamatergic akin to the central synapses found in the vertebrate CNS. The glutamate receptors at the nmj are tetramers made up of four subunits: GluRIIA or GluRIIB, GluRIIC, GluRIID and GluRIIE. Receptors carrying the A or the B type subunit differ in their conductance properties and are shown to be reciprocally regulated such that increase in the levels of GluRIIA leads to corresponding decrease in GluRIIB (Petersen et al., 1997; DiAntonio et al., 1999; Marrus et al., 2004). The mechanisms that regulate glutamate receptor expression, clustering and turn-over are still poorly understood. A classic study by Broadie and Bate more than two decades ago showed that expression of GluRIIA during embryogenesis takes place in two waves. The first wave is autonomous to the muscle and independent of innervation, while the second wave takes place is dependent on innervation. Clustering of the receptors is also dependent on contact between the nerves and muscle (Brodie and Bate, 1993). In the absence of innervation or electrical activity, the muscles fail to upregulate receptor expression in the second wave and clustering fails (Brodie and Bate., 1993). A more recent analysis of the kinetics of GluRIIA transcription suggests that most of the mRNA required for protein synthesis during the embryonic and larval stages is transcribed during embryogenesis with a sharp increase in transcript levels occurring around the time of nerve-muscle contact. An equally sharp decrease in mRNA levels is seen soon after, with no further increase in mRNA levels. The change in GluRIIA protein levels appears to be more gradual and steadily increases through embryonic and larval stages (Ganesan et al., 2009) suggesting that most of the mRNA synthesized during embryogenesis for use during larval stages is sequestered as ribonucleoparticles for subsequent use.

The molecular nature of the pre-synaptic inputs that regulate post-synaptic receptor expression, clustering and maintenance of GluRIIA clusters have remained elusive. Recently, Lola was identified as a transcription factor regulating the expression of many post-synaptic components including GluRIIA. Interestingly Lola activity was found to be and sensitive to neuronal activity such that increased stimulation led to downregulation of Lola activity (Fukui et al., 2012). The molecular identity of other signals, if any, that might function as part of this pre-synaptic regulatory network are not known.

*Drosophila* Mon1(DMon1) is a conserved endocytic factor which in complex with CCZ1 helps recruit Rab7 onto endosomes, thus facilitating the conversion of an early endosome to a late endosome. The protein was first identified in yeast as a factor required for all fusion events to the lysosome (Wang et al., 2002; Wang et al., 2003). This function appears to be conserved across species from yeast to mammals including plants (Poteryaev et al., 2010; Nordmann et al., 2010; Kinchen and Ravichandran., 2010; Yousefian et al., 2013; Cui et al., 2014). While the cellular function of Mon1 is now well established, its physiological role is only beginning to be addressed. For example in Arabidopsis, *mon1* mutants function show poor male fertility due to delayed tapetal degeneration and programmed cell death (Cui et al., 2017). In *Cryptococcus neoformans* Mon1 is essential for virulence (Son et al., 2018). We had previously generated and reported a mutation in *Dmon1* (*Dmon1*^*Δ181*^) generated through a P-element excision that deletes the C-terminal region of DMon1. *Dmon1*^*Δ181*^ mutants show poor viability and motor abilities. The synaptic morphology in these animals is altered. However, a striking phenotype observed in these animals was the elevated levels of GluRIIA in synaptic, and often, the extrasynaptic regions. The requirement of *Dmon1* appears to be primarily neuronal, with neuronal knock-down of *Dmon1* phenocopying the GluRIIA phenotype and expression of *Dmon1* in neurons but not muscle, being able to rescue the mutant defects (Deivasigamani et al., 2015). The mechanism by which DMon1 regulates GluRIIA expression is not clear. Given the relationship between DMon1 and Rab7, we wondered whether Rab7 might play a role in this process. In this study, we have focused our attention to determining whether DMon1 and Rab7 interact to regulate GluRIIA levels at the larval neuromuscular junction.

Here, we first describe a new allele of *Dmon1* previously identified and referred to as *pog*^*1*^ (Matthew et al., 2009). Using genetics and sequencing, we demonstrate that *pog*^*1*^ (hereafter referred to as *Dmon1*^*156*^) is an allele of *Dmon1* with a stop codon at residue 156 of the amino acid sequence. Like the C-terminal deletion mutant, *Dmon1*^*156/Δ*181^ larvae show impaired motor abilities and shortened lifespan. At the synapse these mutants show an increase in GluRIIA levels. To evaluate the role of Rab7 in the regulation of GluRIIA, we knocked down *rab7* in neurons using RNAi. Interestingly, while neuronal knock-down leads to an increase in the intensity of GluRIIA staining, downregulation of *rab7* in the muscle has little effect. Further, there appears to be a dose dependent effect with Rab7^CA^: high levels of expression leads to an increase in GluRIIA. We show that *Dmon1*and *rab7* interact to regulate GluRIIA levels since a transheterozygous mutants carrying one copy each of the *Dmon1*^*Δ181*^ and *rab7*^*1*^ mutation show an increase in GluRIIA levels which are comparable to homozygous *Dmon1*^*Δ181*^. Furthermore, expression of Rab7^CA^ in a *Dmon1*^*Δ181*^ mutant background is able to suppress the GluRIIA phenotype suggesting that the two genes are likely to be part of the same regulatory pathway. Our results thus demonstrate a role for a presynaptic DMon1-Rab7 dependent endosomal pathway in regulating post-synaptic receptor levels.

## Results

### Identification and characterization of *Dmon1*^*156*^

*pog*^*1*^ is an EMS mutant first described as a mutation affecting germband extension during gastrulation in early embryogenesis (Matthew et al., 2009). *pog*^*1*^ is homozygous lethal and the mutants fail to survive beyond the second instar larval stage. It showed non-complementation with *Df(2L)9062* indicating that the mutation is likely to be in either *Dmon1* or *smog*-two genes that are uncovered by the deficiency line. We crossed *pog*^*1*^ to *Dmon1*^*Δ181*^ and *Dmon1*^*Δ129*^ mutants: the former carries a deletion restricted to the 3’ region of *Dmon1*, while the latter has a deletion spanning the 3’ of *Dmon1* and 5’ region of the adjacent *smog* gene (Deivasigamani et al., 2015). Non-complementation was observed in both cases indicating that a mutation is likely to be in *Dmon1*. Indeed, sequence analysis of *Dmon1* in *pog*^*1*^ showed presence of single base pair change leading to an amber mutation at residue 157, suggesting absence of a full-length protein. We hereafter refer to *pog*^*1*^ as *Dmon1*^*156*^. We have previously shown that homozygous *Dmon1*^*Δ181*^ escaper adults exhibit a shortened lifespan and strong motor defects. Consistent with this, both *Dmon1*^*156/Δ181*^ and *Dmon1*^*156*^*/Df(2L)9062* animals were short-lived and showed poor motor abilities. To test if neuronal expression of *UAS-Dmon1:HA* rescues these defects, we expressed the transgene in *Dmon1*^*156/Δ181*^ mutants and found that both defects could be rescued completely to match wildtype animals (Fig.1.B & C).

**Fig. 1.**
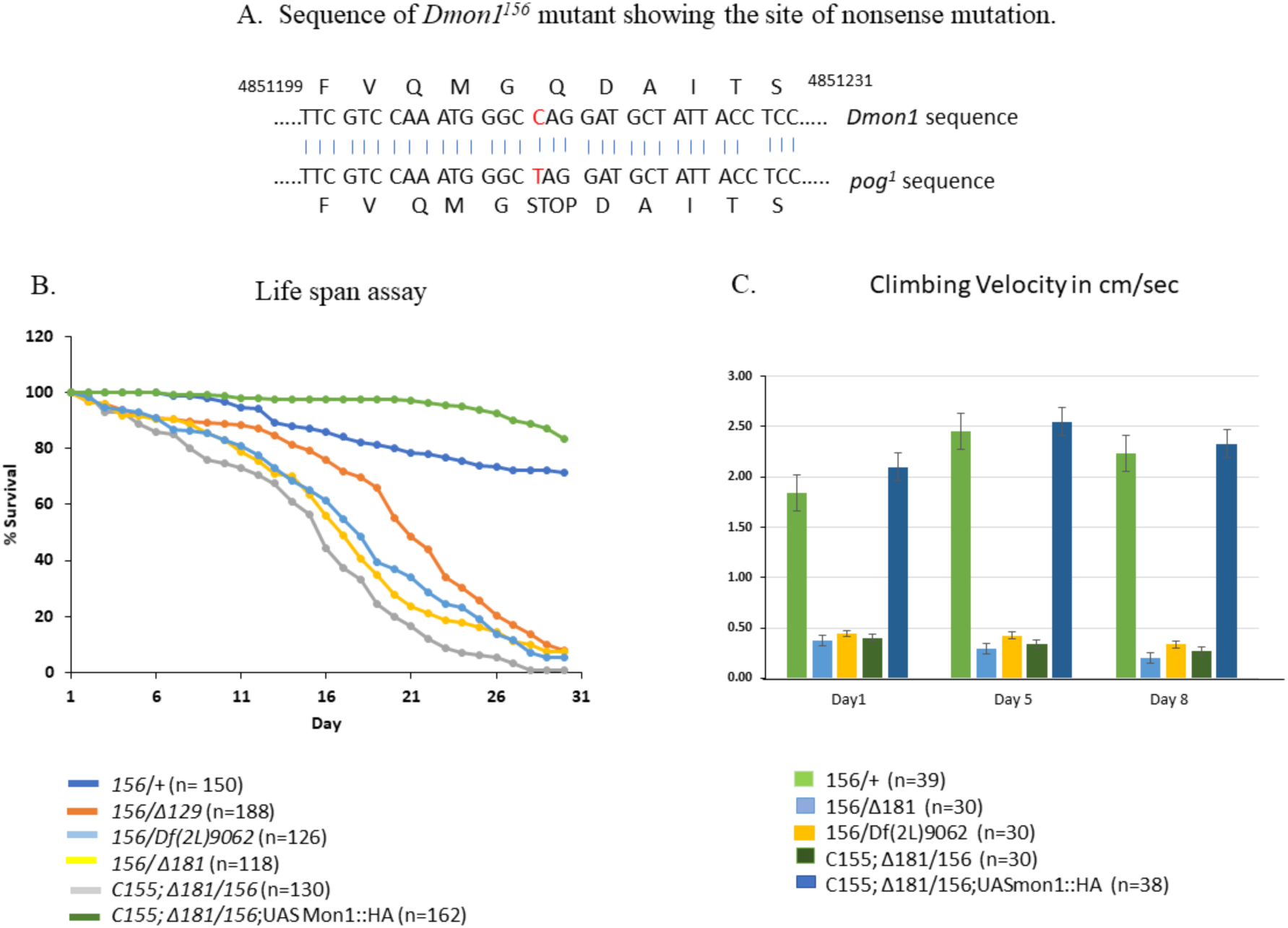
Identification and characterization of *Dmon*^156^ allele. **(A)** Shown is the amber mutation in Dmon1 that generates a stop codon at position 156 of the amino acid sequence of DMon1. **(B)** A comparison of the lifespan of different allelic combinations of *Dmon1*^*156*^ mutants. The strongest allelic combination *Dmon1*^*156/Δ181*^ shows a half-life of approximately 17-18 days. Neuronal expression of *Dmon1: HA* rescues life-span to wild-type levels. **(C)** Shown is a graph of the climbing speed of wildtype and *Dmon1* mutants flies at 1, 5 and 8 days post-eclosion. Motor abilities are compromised in *Dmon1*^*156*^ mutants. This is rescued to wild type levels upon neuronal expression of Mon1: HA. Climbing speed on day 1: Control (*Dmon1*^*156/+*^): 1.84±0.063 versus *Dmon1*^*156/Δ181*^: 0.37±0.033 versus *Dmon1*^*156*^*/Df(2L)9062:* 0.44±0.033 versus C155-GAL4; *Dmon1*^*156/Δ181*^: 0.40±0.027 versus C155-GAL4; *Dmon1*^*156/Δ181*^*>UAS-Mon1: HA:* 2.10±0.146. Climbing speed on day 5: Control (*Dmon1*^*156/+*^): 2.45±0.072 versus *Dmon1*^*156/Δ181*^: 0.29±0.032 versus *Dmon1*^*156*^*/Df(2L)9062:* 0.42±0.036 versus *C155*-GAL4; *Dmon1*^*156/Δ181*^: *0.34*±0.027 versus *C155*-GAL4; *Dmon1*^*156/Δ181*^*>UAS-Mon1: HA: 2.55*±0.198. Climbing speed on day 8: Control (*Dmon1*^*156/+*^): 2.24±0.067 versus *Dmon1*^*156/Δ181*^: 0.20±0.032 versus *Dmon1*^*156*^*/Df(2L)9062:* 0.34±0.032 versus *C155*-GAL4; *Dmon1*^*156/Δ181*^: 0.27±0.024 versus C155-GAL4; *Dmon1*^*156/Δ181*^*>UAS-Mon1: HA: 2.33*±0.138.

Interestingly, the lethality in homozygous *Dmon1*^*156*^ mutants could not be rescued by expression of *UAS-Dmon1:HA* suggesting that there are likely to be other second site ‘hits’ that contribute to lethality in these mutants. However, given the rescue of lethality and climbing defect of *Dmon1*^*156/Δ181*^ by expression of *UAS*-*Dmon1:HA* supports and validates *Dmon1*^*156*^ an allele of *Dmon1*.

### Characterization of Synaptic phenotypes in *Dmon1*^*156*^

*Dmon1*^*Δ181*^ mutant larvae show defects in synaptic morphology: the boutons tend to larger and odd shaped with many supernumerary or satellite boutons (Fig 2B&C; Deivasigamani et al., 2015). A similar phenotype was observed in *Dmon1*^*Δ181/156*^ larvae: the boutons were often bigger and, like the other alleles, showed many more supernumerary boutons (Fig. 2D&I). However, there was no change in bouton number.

**Fig. 2.**
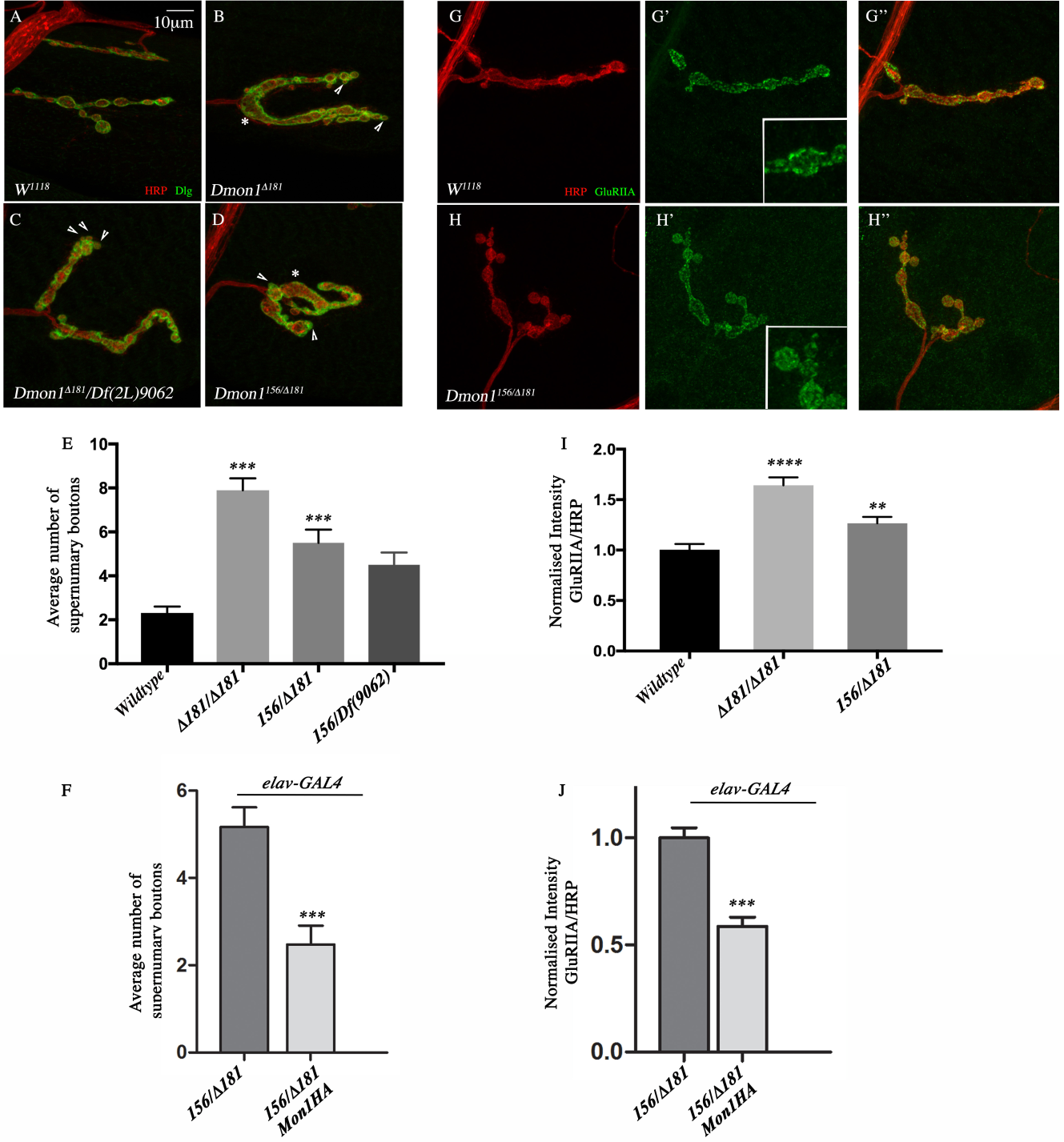
Characterization of the synaptic phenotype in *Dmon1*^*156*^. (A-D) Muscle 4 synapse stained with HRP (red) and Dlg (green). **(A)** Wildtype (*w*^*1118*^) synapse. Note presence of bouton arranged in a ‘pearl on a string’ manner. (B-C) Synapse of a homozygous *Dmon1*^*Δ181*^ **(B)** and *Dmon1*^*Δ181*^/*Df(2L)9062* **(C)** larva. Note presence of fused and irregularly shaped bouton (asterisk) and satellite boutons (arrowheads). **(D)** *Dmon1*^*156/Δ181*^ larvae exhibit a similar synaptic morphology with irregularly shaped boutons and satellite or supernumerary boutons. **(E)** Quantification of the average number of satellite boutons per synapse in different alleles of *Dmon1*. A significant increase in satellite boutons is seen in the mutants (*w*^*1118*^: 2.31±0.286, n=19; *Dmon1*^*Δ181*^: 7.89±0.540, n=19; *Dmon1*^*156/Δ181*^: 5.50±0.600, n=18; *Dmon1*^*156*^*/Df(2L)9062:* 4.50±0.562, n=16). **(F)** The satellite bouton phenotype in *Dmon1*^*156/Δ181*^ is suppressed by expression of DMon1:HA (*elav*-GAL4; *Dmon1*^*156/Δ181*^ :5.16±0.460, n=24, *elav*-GAL4; *Dmon1*^*156/D181*^*>UAS-Mon1:HA* :2.47± 0.430, n=23). (G-H) A muscle 4 synapse from *w*^*1118*^ **(G)** and *Dmon1*^*156/Δ181*^ **(H)** stained with HRP (red) and GluRIIA (green). Note the increase in intensity of GluRIIA. **(I)** Quantification of the intensity of GluRIIA in wildtype (1±0.056, n=16), *Dmon1*^*Δ181*^ (1.744±0.085, n=17) and *Dmon1*^*156/Δ181*^ (1.292±0.065, n=20) animals. The latter show a 30% increase in GluRIIA intensity compared to a near 70% increase in *Dmon1*^*Δ181*^. (J) Expression of DMon1: HA rescues the GluRIIA phenotype in *Dmon1*^*156/Δ181*^ mutants. *** indicates P<0.0001, ** indicates P<0.001 and * indicates P<0.01.

*Dmon1*^Δ181^ mutants show elevated levels of GluRIIA at post-synaptic densities. On an average, the observed increase is nearly two-fold (Deivasigamani et al., 2015). An increase in GluRIIA levels was also observed in *Dmon1*^Δ181/156^ animals (Fig. 2H-H’’). However, unlike *Dmon1*^Δ181^ the increase was approximately 30% (Fig. 2J). The phenotypes associated with synaptic morphology and GluRIIA was suppressed upon expression of Mon1:HA, confirming that the phenotypes are indeed due to loss of *Dmon1* function (Fig. 2K&L). This further validates *Dmon1*^156^ as an allele of *Dmon1* and supports our previous observations on DMon1 being a negative regulator of GluRIIA levels at the larval nmj.

### Loss of Dmon1 does not alter quantal size or quantal content

Overexpression of GluRIIA increases post-synaptic sensitivity leading to increase in quantal size which is the response to release of a single neurotransmitter vesicle. These animals also show an increase in the evoked response or evoked junction potential (EJP) but no change in quantal content or the total number of vesicles released (Petersen et al., 1997).

*Dmon1*^*Δ181*^ mutants show nearly a 2-fold increase in GluRIIA levels. To determine whether this increase in receptor levels has similar physiological effects as GluRIIA overexpression, we carried out intracellular recordings on *Dmon1*^*Δ181*^ and *Dmon1*^Δ181^/*Df(2L)9062* animals. Consistent with the increase in GluRIIA, in both genotypes, a significant increase in EJP was observed (FIG. 3B-C & H) which was rescued by neuronal expression of *UAS-Dmon1*:*HA* (Fig. 3E & H). Surprisingly, and in contrast to overexpression of GluRIIA, we failed to observe any change in quantal size, frequency of the mEPSPs (Fig. 3F-G) or quantal content (Fig. 3I). The lack of an increase in post-synaptic sensitivity suggests that either many of the GluRIIA positive receptors are non-functional or that there is compensation possibly due to the decrease in vesicle size and/or the decrease in GluRIIB which has been observed in these mutants (Deivasigamani et al., 2015).

**Fig. 3.**
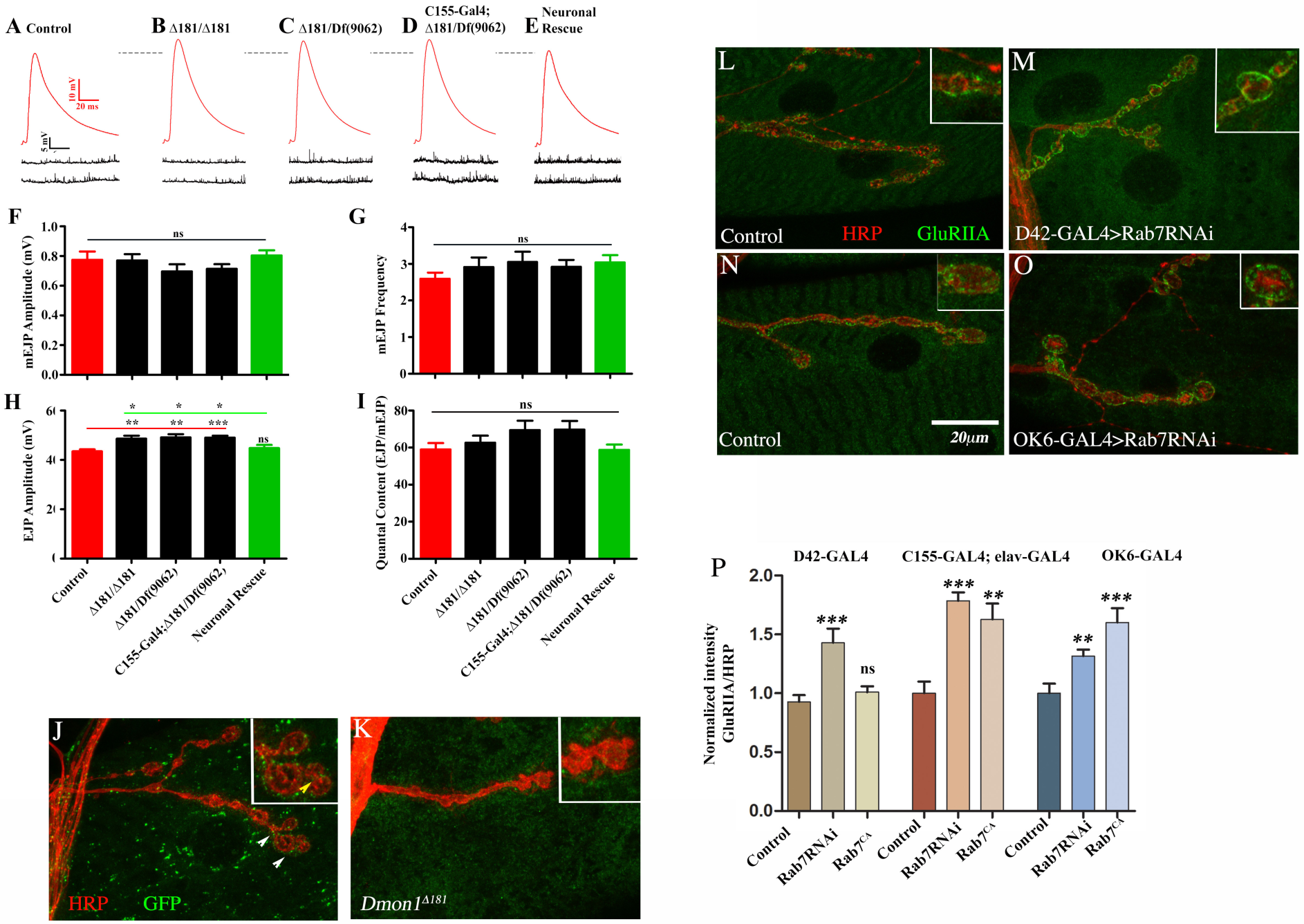
(A-E) Representative traces of evoked (Red) and spontaneous (Black) responses of indicated genotypes. Recordings were carried out in 1.5 mM Ca^2+^ containing HL3 and nerves innervating muscle 6/7 were stimulated at 1 Hz to record evoked responses. **(F)** Histogram showing average mEJP amplitude of Control (0.77±0.05 mV), mutants (*Dmon1*^*Δ181*^, 0.77±0.04 mV; *Dmon1*^*Δ181*^/*Df(2L)9062*, 0.70±0.05 mV), control for rescue (*C155*-GAL4/+; *Dmon1*^*Δ181*^/ *Df(2L)9062*, 0.71±0.03 mV) and rescued animals (0.80 ± 0.03 mV). The numbers in the bars represent number of animals used for recordings and quantification. Error bars represent standard error of the mean (SEM).Statistical analysis based on two-tailed Student’s t-test.^ns^ P>0.5 **(E)**Histogram showing average mEJP frequency of Control (2.59 ± 0.16 Hz), mutants (Δ181/Δ181, 2.91 ± 0.25 Hz; Δ181/Df. 9062, 3.05 ±0.27 Hz), control for rescue (elavC155/+; Δ181/Df. 9062, 2.92 ±0.18 Hz) and rescued animals (3.04 ±0.19 Hz). The numbers in the bars represent number of animals used for recordings and quantification. Error bars represent standard error of the mean (SEM). Statistical analysis based on two-tailed Student’s t-test.^ns^P>0.5 **(G)**Histogram showing average EJP amplitude of control (43.47± 0.08 mV), mutants (Dmon1^Δ181^, 48.64 ± 1.17 mV; Dmon1^Δ181^/*Df(2L) 9062*, 49.12 ± 1.33 mV), control for rescue (*C155*-GAL4; *Dmon1*^*Δ181*^*/Df (2L) 9062*, 49.05 ±0.77 mV) and rescued animals (*C155*-GAL4; *Dmon1*^*Δ181*^*/Df (2L) 9062>UAS-Mon1:HA*, 44.78± 1.31 mV). The EPSP amplitudes reveal significantly enhanced evoked potentials in *Dmon1* alleles, which are rescued significantly by ectopically expressing Mon1 in the mutants. The numbers in the bars represent number of animals used for recordings and quantification. Error bars represent standard error of the mean (SEM). Statistical analysis based on two-tailed Student’s t-test. * P≤0.05, ** P≤0.01. *** P≤0.001 **(H)**Histogram showing average quantal content of Control (58.97± 3.45), mutants (*Dmon1*^*Δ181*^, 62.67± 3.76; *Dmon1*^*Δ181*^/*Df(2L) 9062*, 69.46 ± 5.07), control for rescue (*C155*-GAL4; *Dmon1*^*Δ181*^*/Df (2L) 9062*, 69.70 ± 4.66) and rescued animals (*C155*-GAL4; *Dmon1*^*Δ181*^*/Df (2L) 9062>UAS-Mon1:HA*, 58.76 ± 2.85). The numbers in the bars represent number of animals used for quantification. Error bars represent standard error of the mean (SEM).Statistical analysis based on two-tailed Student’s t-test.^ns^P>0.5 **(I)** Histogram showing average quantal content of control (58.97 ± 3.45), mutants (Δ181/Δ181, 62.67 ± 3.76; Δ181/Df. 9062, 69.46 ± 5.07), control for rescue (elav^C155^/+; Δ181/Df. 9062, 69.70 ± 4.66) and rescued animals (58.76 ± 2.85). The numbers in the bars represent number of animals used for quantification. Error bars represent standard error of the mean (SEM). Quantal content was obtained by dividing average EJP amplitude with average mEJP amplitude of individual recordings. Statistical analysis based on two-tailed Student’s t-test. ^ns^P>0.5 (J-K) Rab7^EYFP^ line stained with anti-HRP and anti-GFP **(J)** Rab7^EYFP^ in *Dmon1*^*Δ181*^ mutants **(K)**. Green puncta in (J) shows localization of Rab7:EYFP at the NMJ. Note the distribution of Rab7 positive endosomes in the muscle. Inset shows presence of puncta in the pre-synaptic compartment and perisynaptic regions. The puncta are completely absent in *Dmon1*^*Δ181*^ mutants. **(L-O)** Representative images of the effect of downregulation of *rab7* on GluRIIA levels using D42-GAL4 (L,M) and OK-GAL4 (N,O) lines. Note the increase in receptor levels (M and O). **(P)** Graph showing the change in staining intensity of GluRIIA in upon expression of *rab7RNAi* and *rab*7^CA^ with different drivers. (D42-GAL4: Control (1±0.056, n=13) versus *rab7RNAi* (1.430±0.119, n=13) versus *rab7*^*CA*^ (1.009±0.049, n=9). *C155*-GAL4; *elav*-GAL4: Control (1±0.099, n=10) versus *rab7RNAi* (1.785±0.072, n=20) versus *rab7*^*CA*^ (1.628±0.134, n=12). OK6-GAL4: Control (1±0.082, n=18) versus RNAi (1.316±0.054, n=20) versus *rab7*^*CA*^ (1.602±0.12, n=19) Note the increase in intensity of GluRIIA in RNAi animals. Expression of *rab* ^*CA*^ also leads to increase in GluRIIA in a dose dependent manner. *** indicates P<0.0001, ** indicates P<0.001 and * indicates P<0.01.

### Neuronal knock-down of Rab7 increases GluRIIA

The above results show that loss of *Dmon1* leading to increase in GluRIIA levels alters neurotransmission. However, the mechanism by which Dmon1 regulates GluRIIA levels is not clear. Given that the conserved function of Mon1 is to recruit Rab7 we sought to determine whether the regulation of GluRIIA is Rab7 dependent. As a first step, we studied the localization of Rab7 at the neuromuscular junction using a genome engineered Rab7^EYFP^ line (Dunst et al., 2015) by staining these larvae with anti-HRP and anti-GFP antibodies. Numerous GFP positive puncta were seen distributed all over the muscle. Interestingly, these puncta were also found near boutons in the peri-synaptic regions (Fig. 3J). Faint GFP positive puncta were also detected inside boutons suggesting presence of late endosomes in the pre-synaptic compartment (Fig. 3J inset). These GFP positive puncta were completely absent in *Dmon1*^*Δ181*^ mutants further reconfirming the role of *Dmon1* in recruiting Rab7 onto vesicles.

Next, we checked whether knock-down of Rab7 in neurons alters GluRIIA levels. Surprisingly, expression of Rab7 RNAi using D42-GAL4 resulted in a significant increase in the intensity of GluRIIA staining (Fig. 3M & P). An approximate increase of 30% was observed with OK6-GAL4 (Fig. 3O &P). A near two-fold increase in staining intensity was observed upon expression of RNAi using a double GAL4 line namely *C155*-GAL4; *ela*v-GAL4 (Fig. 3P). We examined if knock-down of Rab7 in the muscle alters the expression or localization of GluRIIA. Curiously, expression of Rab7RNAi using C57-GAL4 and Mhc-GAL4 had little effect on GluRIIA levels (Supplementary Figure 1).

Based on our results with *Rab7RNAi*, we checked if expression of a constitutively active Rab7 (Rab7^CA^) leads to a decrease in GluRIIA levels. Interestingly, the effect appeared to be dose dependent: while overexpression with *D42*-GAL4 did not result in any significant change in staining intensity, a significant increase in receptor levels was observed with *C155*-GAL4; *elav*-GAL4 and *OK6*-GAL4 lines (Fig. 3P)

### Mon1 and Rab7 interact to regulate GluRIIA

The above results indicate that like *Dmon1*, the *rab7* dependent regulation of GluRIIA is primarily pre-synaptic. To determine whether *Dmon1* and *rab7*^*1*^ interact to regulate GluRIIA, we examined GluRIIA levels in a transheterozygous larvae carrying one mutant copy each of *Dmon1* and *rab7*^*1*^. A significant increase in receptor expression was observed in these animals (Fig4. B). Interestingly, animals heterozygous for *Dmon1*^*Δ181*^ showed a small but consistent increase in synaptic GluRIIA indicating a dose dependent effect (Fig. 4C). In contrast, GluRIIA expression in rab7^1/+^ animals, did not seem to be significantly different from wildtype. The near two-fold increase in GluRIIA intensity in trans-heterozygous *Dmon1*^*Δ181/+*^; *rab7*^*1/+*^ mutants was comparable to the receptor levels observed in *Dmon1*^*Δ181*^, indicating that *Dmon1* and *rab7* interact in a dose dependent manner to regulate GluRIIA. The increase in receptor expression was observed not only at the synapse, but in extrasynaptic regions as well which is often seen in *Dmon1*^*Δ181*^ mutants as well (Fig. 4B). Further supporting these results, a similar synergistic increase in receptor expression was seen in *Dmon1*^*156/+*^; *rab7*^*1/+*^ animals (data not shown). While the steep increase in receptor levels was phenotypically similar to *Dmon1* mutants, the synaptic morphology in these trans-heterozygous animals was very distinct: the boutons in general seemed bigger, well-spaced with fewer satellites (Fig. 4B). To further confirm the genetic interaction between *Dmon1* and *rab7* and to test if they function as part of the same pathway we tested whether expression of Rab7^CA^ suppresses the GluRIIA phenotype in *Dmon1*^*Δ181*^ mutants. Indeed, a strong decrease in GluRIIA staining was observed in these animals (Fig. 4E &F); fewer satellite boutons were observed (Fig. 4G). Together these results indicate that *Dmon1* and *rab7* interact presynaptically to regulate GluRIIA at the larval neuromuscular junction.

**Fig. 4.**
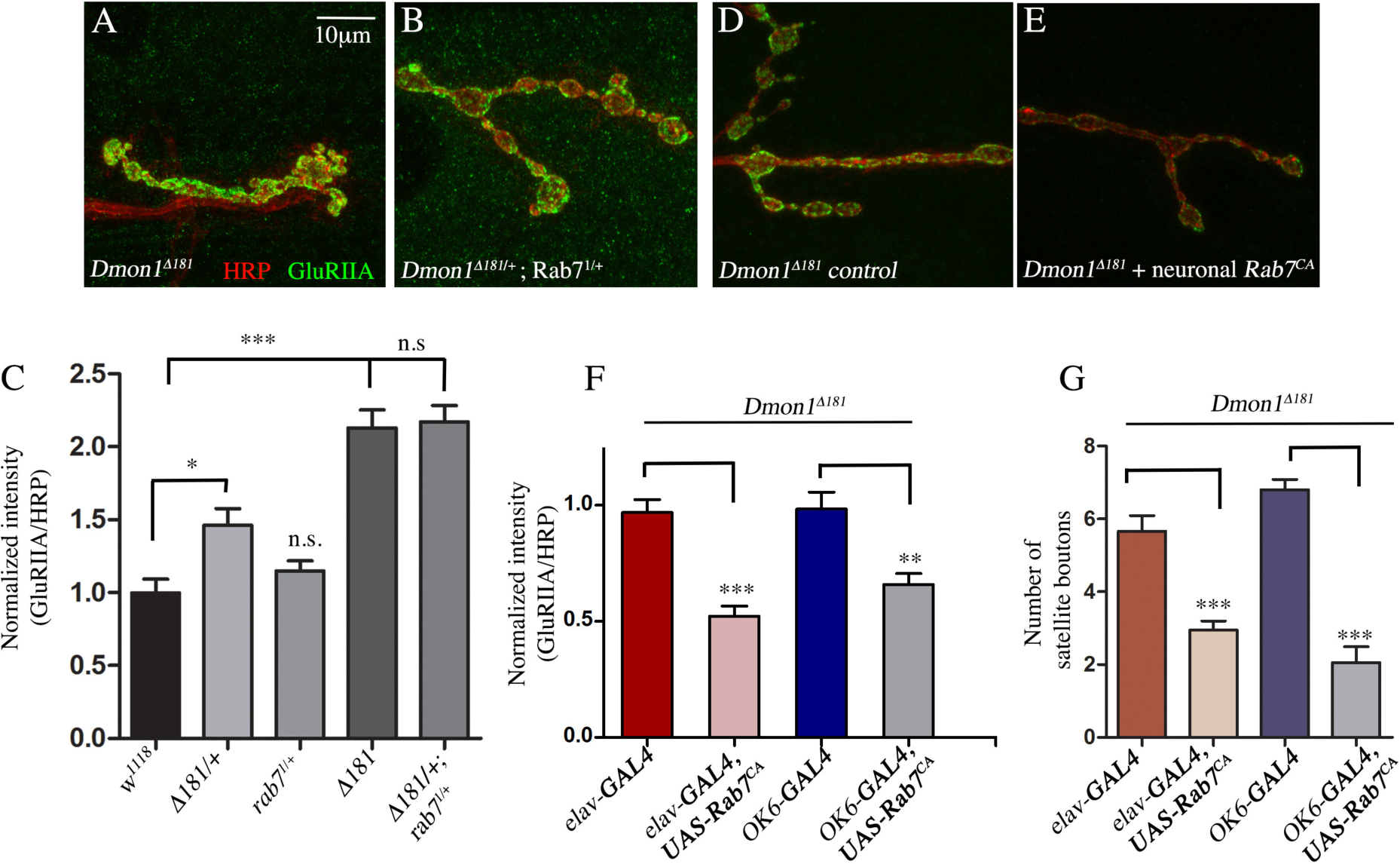
*Dmon1* and *rab7*^*1*^ interact in a dose dependent manner to regulate GluRIIA at the NMJ. **(A-B)** *Dmon1*^*D181*^ (A) and *Dmon1*^*Δ181/+*^; *rab7*^*1/+*^(B) stained with anti-HRP (red) and anti-GluRIIA (green) show comparable levels of GluRIIA. **(C)** Graph showing the normalized intensity values for GluRIIA: HRP ratios. Control (1±0.091, n=6); *Dmon1*^*D181/+*^ (1.461±0.115, n=16); *rab7*^1/+^ (1.149±0.068, n=14); *Dmon1* ^*Δ181*^ (2.129±0.122, n=12); *Dmon1* ^*Δ181/+*^; *rab7*^*1/+*^ (2.172±0.111, n=15). The increase in GluRIIA in transheterozygous mutant animals is comparable to *Dmon1* ^*Δ181*^. **(D-E)** Expression of *rab7*^*CA*^ in *Dmon1* mutants rescues the synaptic and GluRIIA phenotype associated with the mutants. **(F)** Normalized intensity values for GluRIIA: HRP ratios. Mutant control (*Dmon1*^*Δ181*^; *elav*-GAL4/+: 1±0.061, n=18); *Rab7*^*CA*^ ‘rescue’ (*Dmon1*^*Δ181*^; *elav*-GAL4>*UAS-rab7*^*CA*^: 0.515±0.041, n=20); Mutant control (*Dmon1*^*Δ181*^; *OK6*-GAL4/+: 1±0.071, n=20); *rab7*^*CA*^ ‘rescue’ (*Dmon1*^*Δ181*^; *OK6*-GAL4>*UAS-rab7*^*CA*^: 0.685±0.051, n=20). **(G)** Expression of rab7^CA^ rescues synaptic morphology. Fewer satellite boutons are seen. *** indicates P<0.0001, ** indicates P<0.001 and * indicates P<0.01.

## Discussion

Rab proteins play a crucial role in regulating intracellular trafficking by directing endosomes to their appropriate destination, and their dysfunction are associated with various neurodegenerative disease and cancers (Stenmark,H., 2009; Wandinger-Ness and Zerial., 2014; Banworth et al., 2018). Rab7 marks late endosomes and regulates the trafficking and fusion of vesicles with the lysosome. The processes regulated by Rab7 are diverse and range from retrograde trafficking, lysosome positioning, regulation of protein kinases through effectors etc many of which have implications in neurodegenerative disorders (Deinhart et al., 2006; Dodson et al., 2012; Stroupe, C., 2018;). Mutations in Rab7 are associated with Charcot-Marie-Tooth disease 2b (CMT2B) – a type of neurodegenerative disorder that causes axonal degeneration (Verhoeven et al., 2003).

The Mon1-CCZ1 complex functions as a GEF for Rab7, and is essential for its recruitment onto endosomes (Nordmann et al.,2010; Poteryaev et al., 2010). Therefore, it is important to determine the extent to which the phenotypes associated with loss of *mon1* are due to its effect on Rab7. This would be helpful to identify and understand Rab7 independent functions of Mon1 if any. We have addressed this in the context of the *Drosophila* neuromuscular junction where *Dmon1*, through a pre-synaptic mechanism, appears to regulate GluRIIA levels in the muscle.

We find that presynaptic knock-down of *rab7* leads to a significant increase in GluRIIA levels which is comparable to that seen upon knock-down of *Dmon1*. Further, trans-heterozygous mutants with one copy each of *Dmon1*^*Δ181*^ and *rab7*^*1*^ show a dramatic increase in GluRIIA indicating that the two genes interact to regulate GluRIIA. Further, expression of a constitutively active Rab7 (Rab7^CA^) suppresses not only the defects in synaptic morphology but also the increase in GluRIIA seen in *Dmon1* mutants.

Our results indicate that neuronal and not muscle knock-down of *rab7*, leads to an increase in the intensity of GluRIIA staining. This is in variance with a study by Lee and colleagues (Lee et al., 2013) involving knock-down of *tbc1D15-17*-the *Drosophila* ortholog of the mammalian Rab7 GAP, namely *tbc1D15*. RNAi mediated knock-down of *tbc1D15-17* in the muscle but not neuron is seen to affect GluRIIA levels. However, it is to be noted that the effect of *rab7 RNA*i and *rab7*^*CA*^ on GluRIIA has not been evaluated in this study. Moreover, little is known about Rab7 regulation in *Drosophila* and it is possible that the GAP protein regulating Rab7 in the neuron and muscles is different. These questions will need to be addressed in future studies.

The results from overexpression of *rab7*^*CA*^ are a little puzzling (Fig 3). It is not clear why expression with *C155*-GAL4; *elav*-GAL4 leads to an increase in GluRIIA staining. In a recent study Jiminez-Orgaz and colleagues show that TBC1D5 – a GAP for Rab7, in complex with the retromer, controls the localization and cycling of Rab7 such that in the absence of the GAP, the GTP locked form of Rab7 accumulates on the lysosome resulting in defects associated with the non-lysosomal functions of Rab7 (Jimenez-Orgaz et al., 2017). Expression of a constitutively active form of Rab7 showed a similar effect due to accumulation and poor turn-over from the lysosomal membrane due to loss of nucleotide cycling. Given the importance of nucleotide cycling in Rab function, it is possible that expression of Rab7^CA^ beyond a threshold results in sequestration of the activated form on membranes resulting in a *rab7* loss-of-function like phenotype. This will need to tested.

It was a little surprising to see that the increase in *Dmon1*^*156/Δ181*^ is only 30% compared to the average near 1.4 fold increase seen in *Dmon1*^*Δ181/+*^ animals. This would suggest that the increase in staining intensity of GluRIIA in *Dmon1*^*156/Δ181*^ is primarily due to a contribution from a single copy of the *Dmon1*^*Δ181*^ mutation. We do not completely understand the reason for this variation amongst the alleles.

At this point the precise manner by which DMon1-Rab7 axis regulates GluRIIA is not clear. An interesting point to be noted in this context is that DMon1 is released from the pre-synaptic compartment (Deivasigamani et al., 2015). While we cannot completely rule out the possibility of DMon1 and Rab7 having independent effects on GluRIIA, the evidence based on the genetic interaction between the two, and localization of DMon1 suggests that the DMon1-Rab7 complex probably regulates a secretory process that influences GluRIIA levels. At the nmj, signaling molecules such as wingless and synaptotagmin-4 are known to be released via exosomes (Korkut et al., 2009; Korkut et al., 2013) which arise from multivesicular bodies or late endosomes. It would be interesting to test if the regulation of GluRIIA via the DMon1-Rab7 axis involves such a mechanism.

It is intriguing that post-synaptic knock-down of *rab7* does not affect GluRIIA levels in any significant manner. This is somewhat unexpected since in mammalian synapses, endo-lysosomal pathways are known to control the trafficking and turn-over of AMPA receptors (Fernandez-Monreal et al., 2012; Hausser and Schlett., 2017). While the mechanisms regulating GluRIIA turnover at the Drosophila larval nmj are still poorly characterized, the absence of an effect with Rab7RNAi would imply existence of independent pathways. The other possibility could be that knock-down of Rab7 in the muscle is not sufficient enough to elicit a phenotype. These possibilities will need to be explored in greater detail. In summary, our findings here highlight a novel role for the neuronal endo-lysosomal pathway in regulating post-synaptic GluRIIA levels, the details of which will need to be elucidated in future studies.

## Materials and Methods

Flystocks: All stocks were maintained on regular cornmeal agar medium. *pog*^*1*^ (Kind gift from S. Kerridge); *Dmon1*^*Δ181*^ (Deivasigamani et al., 2015); The following stocks were obtained from the Bloomington Stock Centre: UAS-Rab7RNAi (#27051); UAS-Rab7Q67L::YFP (# 9779); UAS-Rab5RNAi (#); UAS-Rab5Q88L (#9774); D42-GAL4 (Rab7 ^EYFP^ (#62545); D42-GAL4 (#8816); OK6-GAL4 (#64199). Rab7^1^ (Kind gift from the Juhász lab; Hegedus et al., 2016.)

### Molecular mapping and behavioral characterization of *Dmon1*^*156*^

*Dmon1*^*156*^, previously referred to as *pog*^*1*^ (Matthew et al., 2009), does not complement *Dmon1*^□*181*^ indicating that the mutation is likely in *Dmon1*. To identify the EMS mutation, we PCR amplified and sequenced the entire *Dmon1* gene from *Dmon1*^*156*^. A single base pair change (C→T) at position 4851219 of the flybase sequence was results in an amber mutation (CAG to TAG) at position 157 of the amino acid sequence of DMon1. The lifespan and motor assays to characterize *Dmon1*^*156*^ phenotype was carried out as described in Deivasigamani et al., 2015.

### Immunohistochemistry

Larval fillets were fixed with Bouins for 15 minutes at room temperature (23-24°C) and stained using standard protocols (Patel et al., 1994). 2% BSA was used for blocking. Anti-HRP (Sigma, 1:1000); anti-GluRIIA (concentrate; DSHB, Iowa state, 1:200); anti-GFP (Thermo Fisher, 1:1000). Imaging was done using a Leica SP8 confocal system. All images were captured with a 63X, 1.4 N.A. objective. Image analysis was carried out using ImageJ (NIH) or FIJI software. Intensity measurements for GluRIIA were carried out as described in *Menon et al*., 2004 and *Deivasigamani et al.*, 2015.

### Electrophysiology

Intracellular electrophysiology recordings were performed as described previously (S.D Chodhury et.al., 2016). Briefly third instar wandering larvae were dissected in modified HL3 saline containing 70 mM NaCl, 5 mM KCl, 20 mM MgCl2, 10 mM NaHCO3, 115 mM sucrose, 5 mM trehalose, 5 mM 4-(2-hydroxyethyl)-1-piperazineethanesulfonic acid (HEPES), and 1 mM EGTA at pH 7.2.For recordings the EGTA was replaced with 1.5 mM CaCl2 in modified HL3 saline (Verstreken et al. 2002). mEPSPs were recorded for 60 S in absence of any stimulation. For evoked responses (EPSPs), motor axons were stimulated at 1Hz and responses were recorded for 1minute. All the recordings were made using sharp glass microelectrodes with 15-25 MΩ resistance from muscle 6 of A2 hemi-segment. Data was analysed using an offline software minianalysis (Synaptosoft)

### Statistical Analysis

Analysis was done using GraphPad Prism software. Two-tailed student’s t-test and ANOVA was used for analysis. All values represented in the figures are mean ± s.e.m.

## Acknowledgements

The authors thank the Bloomington stock centre, Indiana and Developmental Studies Hybridoma Bank (DSHB), Iowa, for fly stocks and antibodies. This work was supported by funds from DBT (BT/PR23318/BRB/10/1597/2017) to AR and CSIR-Senior Research Fellowship to AB.

## Figure Legends

**Supplementary Figure 1.**
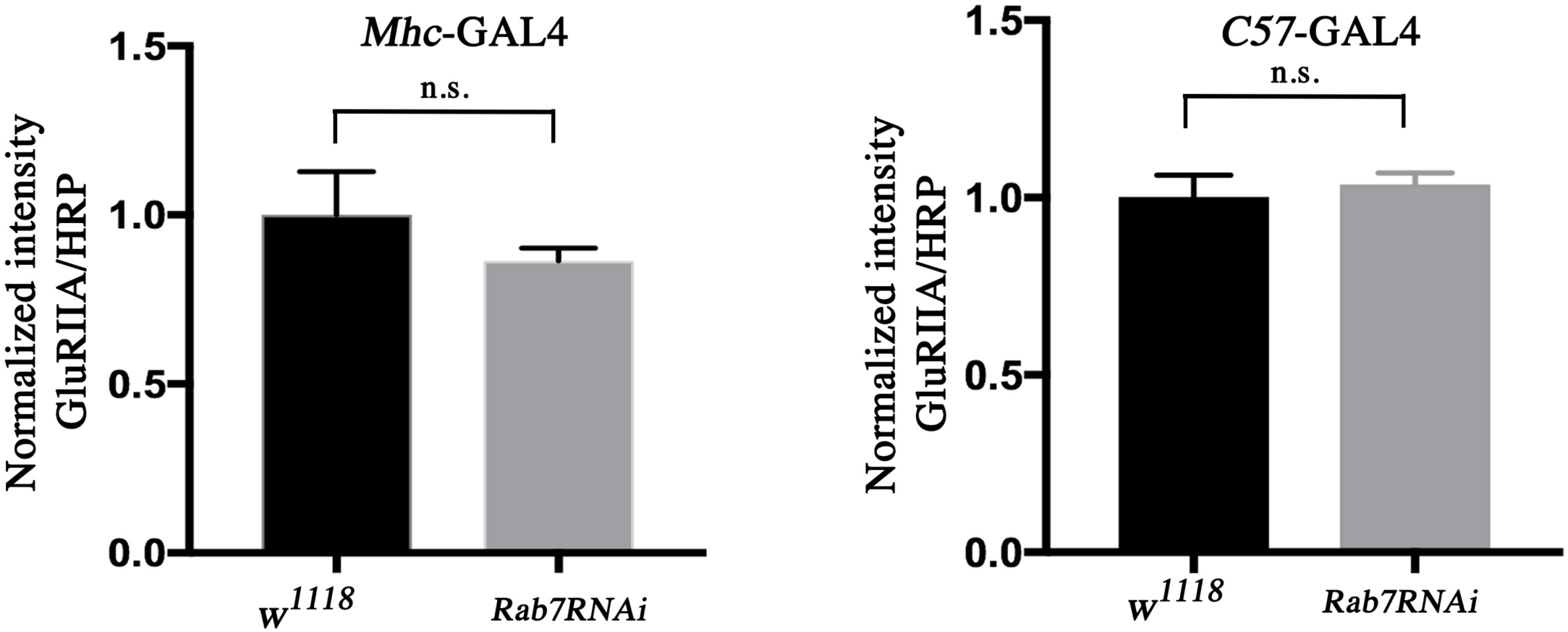
Knock-down of *rab7* in muscles does not alter GluRIIA levels. Expression of UAS-*rab7RNAi* using Mhc-GAL4 and C57-GAL4 does not significantly alter GluRIIA levels at the nmj.

